# A Robust Statistical Approach for Finding Informative Spatially Associated Pathways

**DOI:** 10.1101/2024.03.31.587469

**Authors:** Leqi Tian, Jiashun Xiao, Tianwei Yu

## Abstract

Spatial transcriptomics offers insights into functional localization of cells by mapping gene expression to spatial locations. Traditional focus on selecting spatially variable genes often misses the complexity of biological pathways and biological network dynamics. We introduce a novel framework that shifts the focus towards identifying functional pathways associated with spatial variability, by adapting the Brownian distance covariance test to explore the heterogeneity of biological functions over space. The statistical approach is free of parameter selection. It allows for a deeper understanding of how cells coordinate their activities across different spatial domains through biological processes. By analyzing real human and mouse datasets, the method found significant pathways that were associated with spatial variation, as well as different pathway patterns among inner- and edge-cancer regions. This innovative framework offers a new perspective on analyzing spatial transcriptomic data, contributing to our understanding of tissue architecture and disease pathology.

## Introduction

The rapid developments in Spatially Resolved Transcriptomics (SRT) technology have provided genomic research an unprecedented perspective. The technology can simultaneously capture the expression measurements of thousands of genes at multiple samples and spatial points along with their spatial coordinates within tissues. This combination effectively bridged the gap between transcriptomics and histological topology (1, 2). Technology like Visium integrates microfluidic partitioning with barcoded bead arrays to feature about 5,000 spots with a diameter of 55 μm each, offering a detailed glimpse into the spatial variation of gene expression across whole-tissue sections. And Slide-seq transfers RNA from tissue sections onto a surface arrayed with DNA-barcoded beads measuring 10 μm, enabling the sequencing of RNA from thousands of spatially resolved points within the tissue (3). High-Definition Spatial Transcriptomics (HDST) elevates this further, providing subcellular insights that approach the single-cell level, capturing the heterogeneity of tissue architecture and the nuanced shifts in gene expression patterns with remarkable clarity (4).

Based on the spatial transcriptomics data, researchers can utilize tissue spatial location information when studying gene expression patterns (3). Combined with the use of spatial coordinates, researchers can now explore how gene expression adjusts to dynamic changes in the cellular microenvironment, enabling an unprecedented level of analysis. When analyzing gene expression patterns, identifying highly variable genes (HVGs)(5) is a straightforward approach that has been widely used in single-cell RNA-seq (scRNA-seq) data analysis. After incorporating spatial information, spatially variable genes (SVGs) are defined as genes showing differential expression in different tissue regions or cell types, a definition that not only demonstrates their involvement in various biological processes but also reveals tissue. The heterogeneity among them provides extremely valuable information for biological research (2). Numerous methodologies for detecting spatially variable genes (SVGs) have been introduced in contemporary research, signifying the progression in spatial transcriptomics. SpatialDE distinguishes gene variations as spatial or non-spatial by fitting each gene with a Gaussian process model (6). nnSVG estimates each gene by employing nearest neighbor Gaussian processes within the spatial covariance function (7). SPARK-X, leveraging a non-parametric approach, assesses the covariance similarity between the expression matrix of each gene and the spatial coordinates matrix to conduct hypothesis testing (8). The BSP (big-small patch) method identifies SVGs by comparing variances in gene expression across two spatial granularities (9). Furthermore, an array of methods, including Trendsceek (10) and MERINGUE (11), among others, expands this repertoire.

While gene-centric analyses shed light on the molecular mechanisms linked to specific cellular states, they frequently fall short in grasping the intricacy of biological pathways and network dynamics. The spatial patterning of gene expression transcends random occurrence, mirroring the deep-seated biological processes and interactions. Therefore, our approach strives to move beyond individual gene analysis, advancing towards a more integrated gene network perspective to decipher the biological heterogeneity among tissues. Prior work has already harnessed pathway information within model designs. PASNet incorporates pathway information into a sparse deep neural network to predict long-term survival in glioblastoma multiforme (GBM) patients (12), while PathCNN applies convolutional neural networks to biological pathway images for prognostication in GBM (13). Skillfully integrating biological pathway data not only improves model performance but also enhances the biological interpretability of the models.

Therefore, we introduce a novel method that emphasizes identifying functional pathways/biological processes associated with spatial variability. By concentrating on pathways, our approach delves deeper into the spatial heterogeneity of biological functions and uncovers how cells coordinate their activities through biological processes across different spatial domains. We employ the Brownian distance covariance test to evaluate the dependency of two random vectors with different dimensionality. The test can find informative spatially varying pathways from two aspects: (i) find pathways generally related to spatial location of a tissue, (ii) evaluate the functional difference between pre-specified spatial areas, such as the edge spots and the inner spots within the same region of the tissue.

## Methods

For spatial transcriptomics data, we incorporate both gene expression counts and spatial positioning information. Genes expressed in fewer than a pre-specified proportion of cells (twenty cells in this study) within the raw expression matrix are excluded. Subsequently, the expression counts are normalized and log-transformed using the SCANPY package (14). We define the processed gene expression matrix as *G* ∈ ℝ^*n×r*^ and spatial location matrix as *L* ∈ ℝ^*n×*2^.

### Brownian Distance Covariance Test

The concept of distance covariance (dCov) was introduced as a metric to measure the dependence between two random vectors *X* and *Y* of arbitrary dimensions (15). Compared to Pearson correlation, distance correlation offers an advantage by detecting both linear and nonlinear associations, as well as the flexibility in dimensionality. Consider two random vectors *X* ∈ ℝ^*p*^ and *Y* ∈ ℝ^*q*^, where *p* and *q* are positive integers. The characteristic functions of *X* and *Y* are represented as *f*_*X*_ and *f*_*Y*_, with their joint characteristic function denoted by *f*_*X,Y*_. The discrepancy between between *f*_*X,Y*_ and *f*_*X*_ *f*_*Y*_ can be quantified by the distance ‖*f*_*X,Y*_ −*f*_*X*_ *f*_*Y*_ ‖. Thus, we can test the independence using the hypothesis as the following,

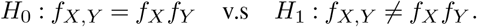

The Distance covariance *V* is designed as a measure of dependence which is defined by

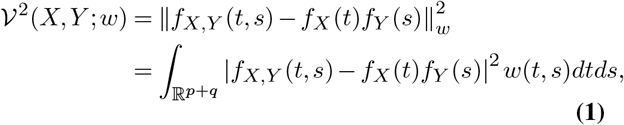

with weight function *w*(*t, s*). The Distance covariance *V* has an important property that *V* ^2^(*X, Y* ; *w*) = 0 of and only if *X* and *Y* are independent. The empirical version of covariance *V* is applied to test the hypothesis. Specifically, with observations 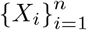 and 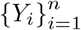, the empirical distance covariance is developed as the square root of

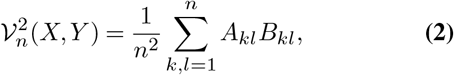

where matrices *A* and *B* are determined by

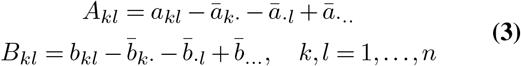

with *a* and *b* which represent the pairwise Euclidean distance matrices for the sets 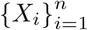 and 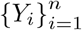 and

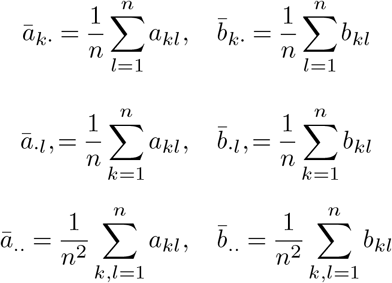

The empirical distance correlation (dCor) is defined as the square root of

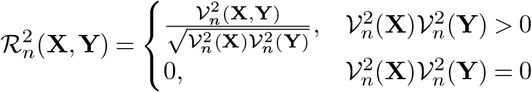

and the p-value for the dCor association is derived via a permutation test (16).

### Testing between pathway gene expression and location

We devised a methodology based on the Brownian distance covariance test to explore the relationship between gene expression relevant to biological processes and spatial location. All involved processes are tested, and the ones with high dCor and significant p-values are recognized to be associated with spatial variation. Such analysis aids in our comprehension of regional variations in tissue functionality from a biological standpoint. Figure 1 depicts the workflow and examples.

**Fig. 1.**
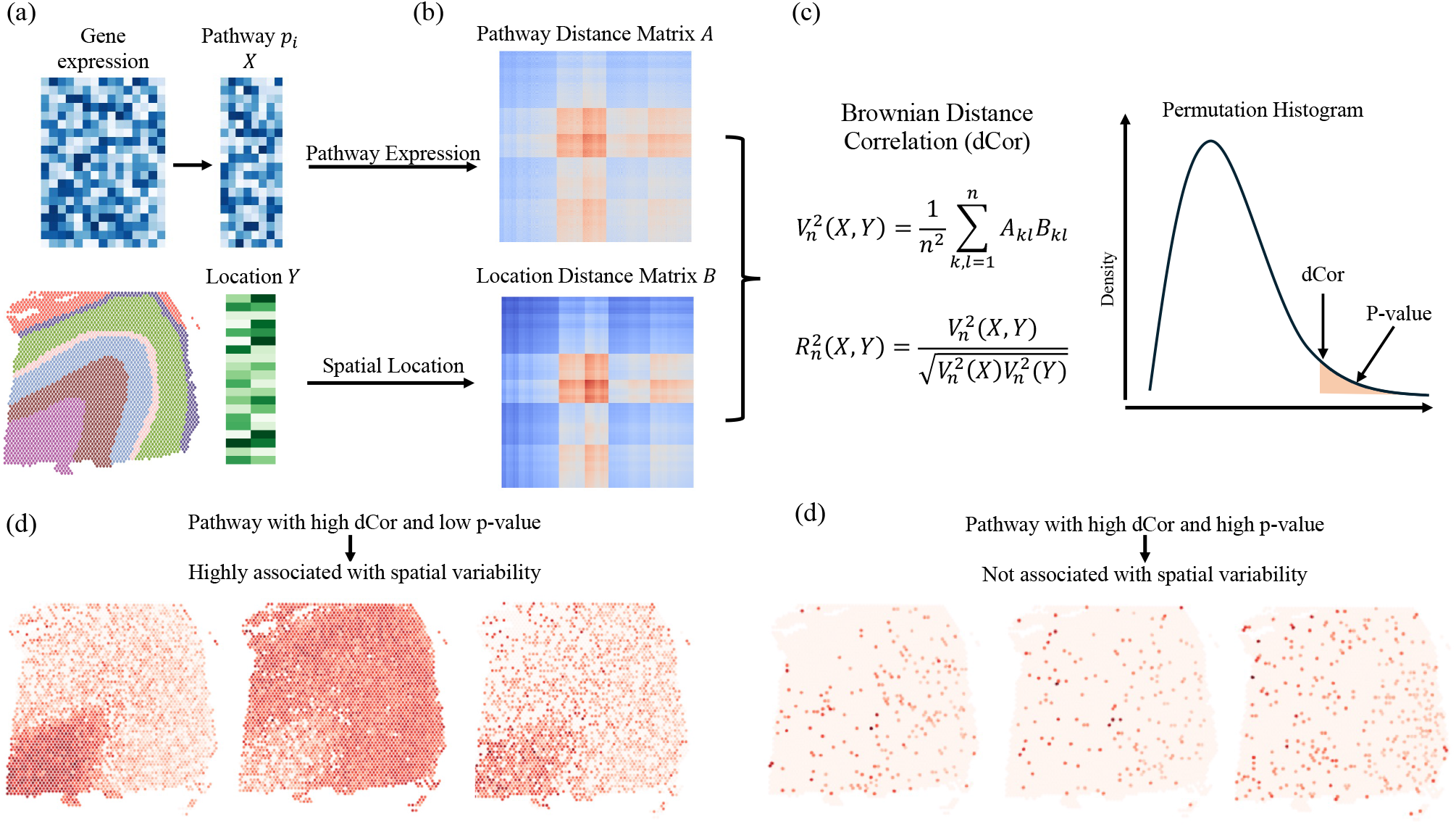
Schematic overview of analyzing the association between pathway expression and spatial location. (a) For a spatial transcriptomic data, the matrix X contains genes involved in pathway *p*_*i*_, and the location matrix *Y* is the spatial coordinate. (b) The Pathway distance matrix *A* and location distance matrix *B* are calculated based on matrix *X* and *Y*. (c) The brownian distance covariance *V*_*n*_(*X, Y*) and brownian distance correlation *R*_*n*_(*X, Y*) are estimated using matrix *A* and *B*. The p-value are estimated by permutation test. (d) Example of heatmap of genes in pathway with high dCor and significant p-value, indicating the pathway is highly associated with spatial variability. (d) Example of heatmap of genes in pathway with low dCor and high p-value, indicating the pathway is not associated with spatial variability.

Give a spatial transcriptomic data with gene expression matrix *G* and location coordinate matrix *Y*, our initial step is pathway information retrieval. We utilize the Gene Ontology (GO) database (17), a comprehensive resource that annotates genes with their associated biological processes. From this database, we focus on biological processes that encompass total gene count ranging from twelve to five hundred. Suppose the filtration results in *m* biological processes, we then denote them as *P* = (*P*_1_, *P*_2_, …, *P*_*m*_). For any *i* ∈ [1, 2, …, *m*], we denote the process *P*_*i*_ encompasses *r*_*i*_ genes, and the list of associated gene names is represented as 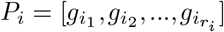 With the gene list *P*_*i*_, we define 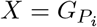 as a sub-matrix of the gene expression matrix *G*, containing only the genes associated with the *i*-th biological process *P*_*i*_ shown in Figure 1(a).

Given the sub-matrix 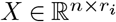, corresponding to pathway *f*_*i*_, and the spatial location matrix *Y* ∈ ℝ^*n×*2^, we compute the pathway distance matrix *A* and the spatial distance matrix *B*, both of which are of dimension ℝ^*n×n*^. These matrices represent the estimated spot-wise distances, with *A* derived from gene expression data and *B* reflecting the distances between spatial coordinates, respectively, shown in Figure 1(b).

Subsequently, we utilize the Brownian distance correlation score to evaluate the spatial association, as depicted in the formula within Figure 1(c). The p-value is determined through a permutation test to assess the statistical significance of this association. After examining all the *m* pathways, the highest test statistics are selected. The biological significance of these pathways is further validated by a comparative review of known tissue-specific functions and structures from the literature. Figure 1(d) illustrates an example heatmap of a pathway characterized by a high distance correlation (dCor) and a significant p-value, suggesting a strong correlation with spatial variability. Conversely, Figure 1(e) presents an opposite scenario. It is evident that one exhibits a distinct spatial pattern, whereas the other appears to follow a random distribution, which is consistent with the test results.

To further explore the spatial organization of gene expression within the highest-ranked pathways, we implement K-means clustering on the gene expression data of these pathways. This approach allows us to visualize and assess the spatial distribution of gene expressions associated with selected pathways. As a comparative benchmark, we conduct a parallel analysis using Highly Variable Genes (HVGs) selected to match the gene count of the analyzed pathways. The effectiveness and robustness of the clustering results are evaluated using the Adjusted Rank Index (ARI), which provides a quantitative measure of clustering quality. The result is shown in the Supplementary Supplemental Figure 1.

### Functional analysis of different regions within the same cell type

To advance our understanding of gene expression pathway dynamics within diverse microenvironments, we advocate for a methodological refinement focused on analyzing the biological functional disparities among different regions within the same cell type. Specifically, we divide cell-type spots into two distinct categories, edge and inner, utilizing available annotation and location information. This categorization facilitates a more comprehensive examination of functional variations, challenging the conventional notion of homogeneity based on cell type similarity. The Brownian distance covariance is a particularly valuable metric for measuring the difference due to its versatility in accommodating random vectors across arbitrary dimensions and its non-negative properties. The dCor values close to zero indicate statistical independence. Figure 2 illustrates the analysis procedure and example.

**Fig. 2.**
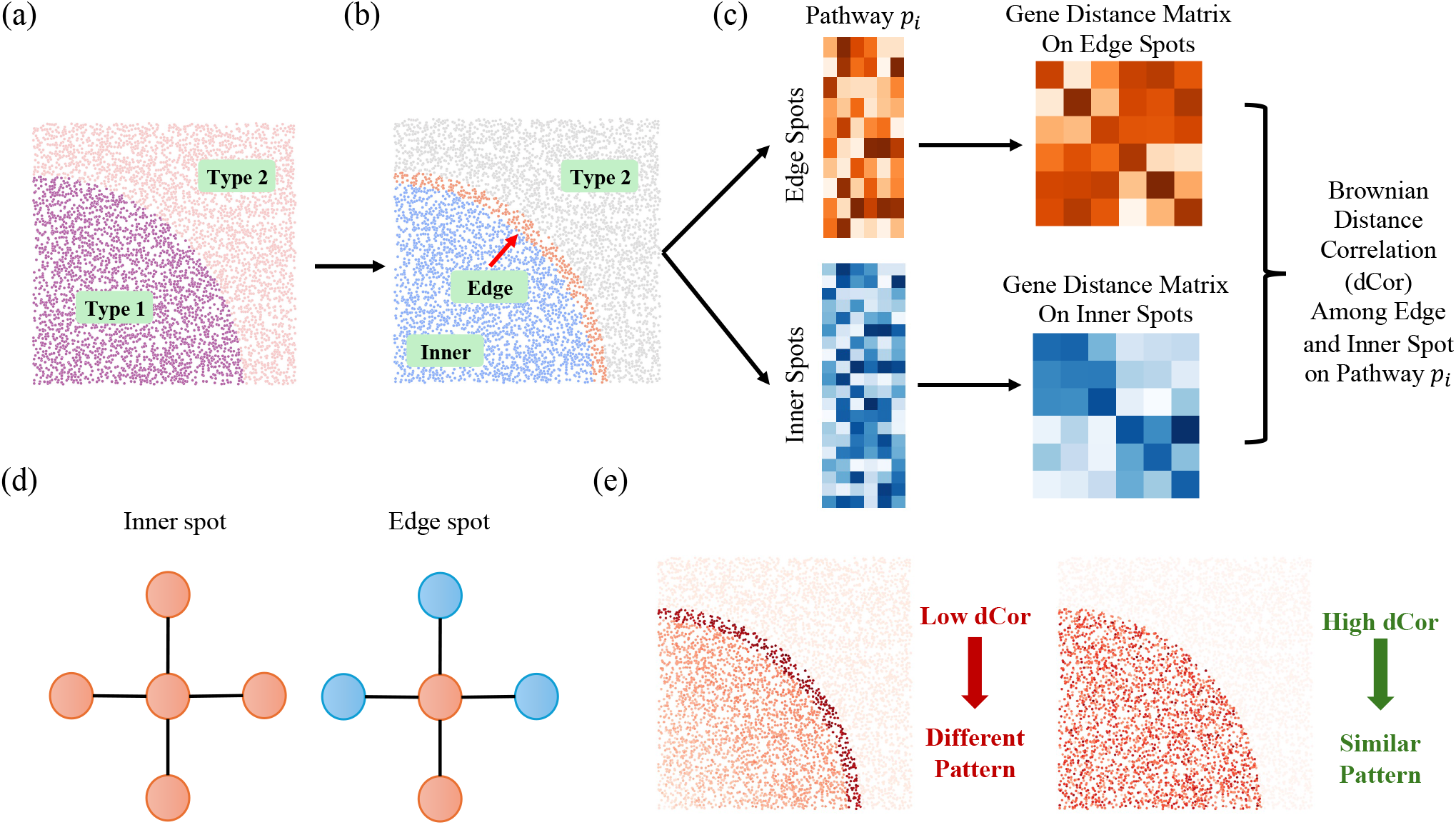
Framework for examining functional variability within the same cell types across different regions. (a) Illustration depicting a spatial microenvironment containing Type 1 and Type 2 cell regions. (b) Focus on Type 1 cell regions identifies distinct areas as either edge or inner regions. Inner spots are represented by blue nodes, edge spots by orange nodes, and Type 2 cell regions by grey nodes. (c) For pathway *p*_*i*_, two sub-expression matrices are delineated; one representing gene expressions at edge spots (orange) and the other at inner spots (blue). Subsequently, gene distance matrices for edge and inner spots are computed separately, facilitating the estimation of distance correlation (dCor). (d) Illustration of inner spot and edge spot. A spot is classified as an inner spot if all neighboring spots belong to the same cell type; otherwise, it is considered an edge spot. (e) Example of different pathway pattern and similar pathway pattern between inner and edge regions.

In datasets annotated with cell type information, we introduce a classification system based on spot location—either inner or edge—determined by the congruence of cell types between a spot and its neighboring regions. A spot is designated as “inner” if its cell type aligns with that of its immediate neighbors; conversely, it is categorized as an “edge” spot if there is a disparity in cell types.

For a given pathway *p*_*i*_, we commence by selecting relevant genes and constructing the expression matrix for our region of interest regarding pathway *p*_*i*_. Genes expressed in fewer than twenty spots within this region are excluded to refine the analysis. The resulting expression matrix, denoted as *X*′ and represented as 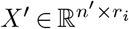, is divided into matrices for edge and inner expressions, symbolized by 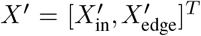with

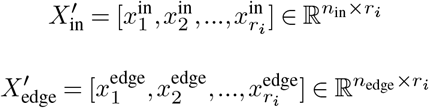

where *n*_edge_ and *n*_in_ denote the number of cells in the edge and inner areas, while *r*_*i*_ signifies the number of genes within the pathway. We calculate the gene Euclidean distance matrices 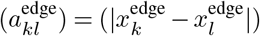 and gene distance matrix on inner spots 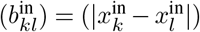, *k, l* = 1, 2, …, *r*_*i*_. According to the formula 3, 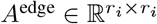 and 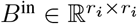 can be calculated and thus we have the dCov

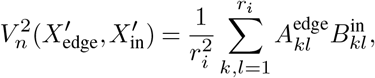

and the dCor 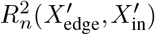 depicting the difference among the two region in pathway *f*_*i*_. Upon evaluating all pathways, we identify those pathways where correlation values are closest to zero. This highlights pathways whose expression in the inner region is statistically independent from their expression in the edge region, thereby shedding light on diverse expression patterns between inner and edge spots.

## Results

### Pathways related to spatial location

#### Human Pancreatic Ductal Adenocarcinoma Study

We applied our method on human pancreatic ductal adenocarcinoma (PDAC) data (18) with 10,000 permutations. Based on histological annotations, the data were categorized into several tissue regions, including cancer, duct epithelium, pancreatic, and stroma, as depicted in Figure 4(a). Our analysis identified multiple significant pathways associated with spatial location variability, as shown in Table 1 which lists each pathway name, the count of related genes, the statistic (dCor), and P-value derived from the our test.

**Table 1.** Most significant pathways associated with spatial variation on PDAC dataset.

| GOID | Term | n_gene | dCor | P-value |
| --- | --- | --- | --- | --- |
| GO:0045104 | intermediate filament cytoskeleton organization | 25 | 0.599 | <0.0001 |
| GO:0030307 | positive regulation of cell growth | 66 | 0.591 | <0.0001 |
| GO:0008544 | epidermis development | 104 | 0.585 | <0.0001 |
| GO:0006959 | humoral immune response | 84 | 0.566 | <0.0001 |
| GO:0010927 | cellular component assembly involved in morphogenesis | 36 | 0.563 | <0.0001 |
| GO:0042692 | muscle cell differentiation | 142 | 0.552 | <0.0001 |
| GO:0010876 | lipid localization | 158 | 0.543 | <0.0001 |
| GO:0048144 | fibroblast proliferation | 53 | 0.542 | <0.0001 |
| GO:0050878 | regulation of body fluid levels | 150 | 0.542 | <0.0001 |
| GO:0010951 | negative regulation of endopeptidase activity | 96 | 0.538 | <0.0001 |

One of the most significant pathways is the organization of the intermediate filament cytoskeleton. Intermediate filament plays a crucial role in providing essential structural support and contributing to cell integrity and signaling within the cytoskeleton. The fundamental process of cancer cell migration depends on the dynamic interplay between the cytoskeleton and cell surface receptors (19). Simultaneously, tumor cells undergo metabolic adaptations to facilitate proliferation and metastasis, requiring the positive regulation of cell growth and cellular tissue architecture to undergo restructuring. Epithelial-to-mesenchymal transition induces a shift from an epithelial to a motile fibroblast-like morphology, which is pivotal within the tumor microenvironment of pancreatic ductal adenocarcinoma (20). This transition has been demonstrated to play a significant role in the invasion and metastasis of PDAC (21). Several other pathways involved in cell growth, differentiation and development, well known for their association with cancer, are also found among the top 10 list, including positive regulation of cell growth, epidermis development, muscle cell differentiation *etc*. Another interesting biological process is humoral immune response. PDAC has been studied for tumor-stroma interaction and cancer microenvironment (22).

#### Mouse Olfactory Bulb Study

In this study, we applied our method to Visium spatial transcriptomic data from the mouse olfactory bulb (MOB) (GSM4656181). Figure 3(a) displays the histological image of the dataset. Using annotations from other histologically marked images, we broadly outlined regions within the image, including the Rostral Migratory Stream (RMS), Granule Cell Layer (GCL), Inner Plexiform Layer (IPL), External Plexiform Layer (EPL), Glomerular Layer (GL), and Outer Nuclear Layer (ONL). Figure 3(b) outlines the significant pathways associated with spatial variation identified by our method. These terms can be broadly categorized into pathways related to steroidogenesis, axonal structure and function, exploratory behavior, and other no-table pathways.

**Fig. 3.**
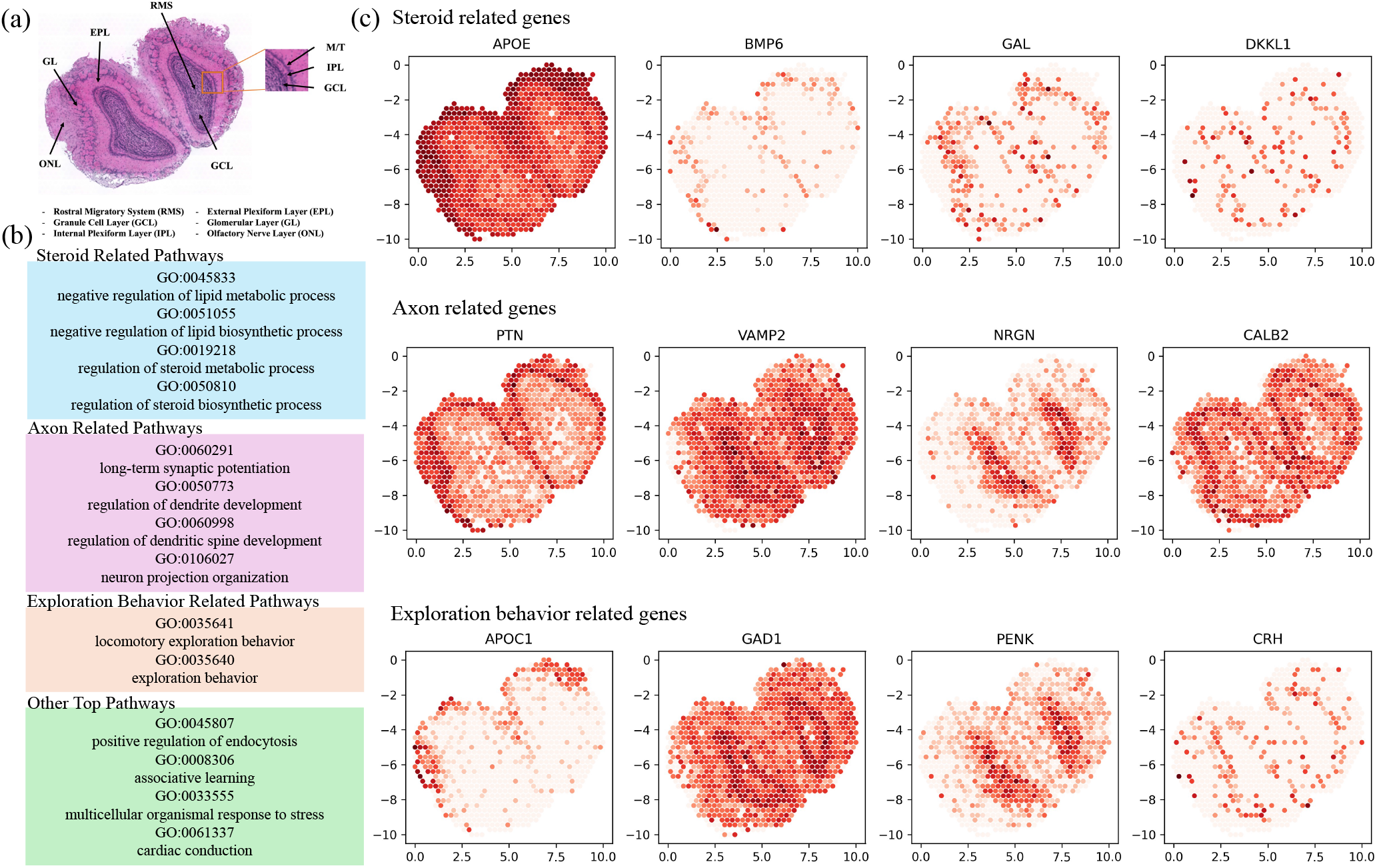
Application of STPath to mouse olfactory bulb (MOB) data. (a) Image of H$E-stained mouse olfactory bulb. (b) Most significant pathways associated with spatial variation identified by STPath.We divide the pathways into four broad categories, steroid related, axon related, exploration behavior related and other pathways. (c) Heatmaps of genes to top pathways related to steroid, axon and exploration behavior.

**Fig. 4.**
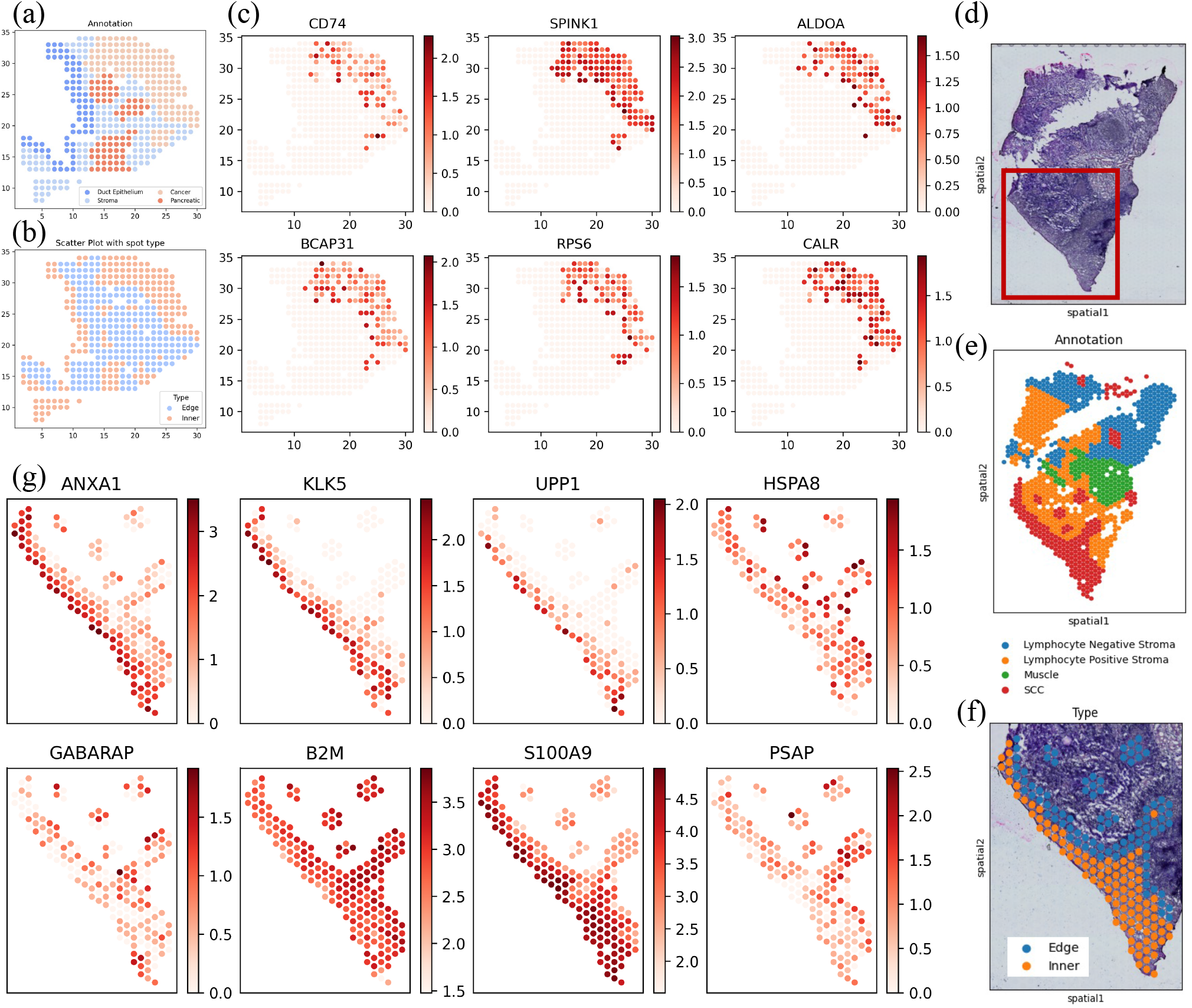
Functional analysis between inner and edge cancer spots on PDAC and OSCC datasets. (a) Annotated regions of PDAC: cancer, pancreatic, ductal and stroma. (b) Depiction of inner and edge spots. (c) Heatmaps of genes related to GO:0042742, GO:0007276. (d) Image of H&E-stained oral squamous cell carcinoma (OSCC). (e) Annotated regions of OSCC: SCC, muscle, lymphocyte negative stroma and lymphocyte positive stroma. (f) Depiction of inner and edge cancer spot of our interest cancer area. (g) Heatmaps of genes related to the most different pathways.

One of the most different parts is lipids-related pathways, including negative regulation of lipid metabolic process (GO:0045833), negative regulation of lipid biosynthetic process (GO:0051055), regulation of steroid metabolic process (GO:0019218), and regulation of biosynthetic process (GO:0050810). Lipids are crucial components of brain composition and function, playing an essential role in maintaining neuronal membrane structural integrity, which is vital for the normal functioning of neurons and the precise transmission of neural signals. The External Plexiform Layer (EPL) primarily consists of dendrites from mitral cells and axons from peripheral neurons. At the same time, the Glomerular Layer (GL) is enriched with synaptic connections between olfactory neurons and secondary neurons (23, 24). The olfactory nerve layer contains olfactory nerve fibers extending from the nasal cavity, involved in the initial reception of olfactory signals (25). Here, the regulation of lipid metabolism is particularly crucial for neuroprotection and preventing cellular damage caused by excessive lipid accumulation. Meanwhile, steroids, including sex hormones and corticosteroids, play critical roles in neuronal development, maintenance, and modulation (26). The regulatory dynamics of steroid metabolism can fine-tune olfactory signaling, affecting stress responses and emotional regulation in the External Plexiform Layer. Studies suggest that steroid medications can improve olfactory nerve regeneration and reduce inflammation at injury sites (27).

Another set of essential terms identified relates to synaptic function and neuronal development. Long-term potentiation (LTP) is a form of enhanced communication between neurons closely associated with learning and memory. GO:0050773 and GO:0060998 refer to the regulation of dendrites and dendritic spines, respectively. These structures are essential for receiving input signals and forming synapses with other neurons. They participate in integrating and regulating signal output in the nervous system (28). Neuronal Projection Organization (GO:0106027) involves the extension and branching of neuronal axons, facilitating communication within the olfactory bulb and potentially influencing the pathways of olfactory information transmission (29). In Figure 3(c), we observed differential gene expression of these terms between the GCL and GL layers. The GCL, despite lacking synapses, plays a central role as an intermediary neuron in olfactory signal processing (30), with inputs from olfactory receptor neurons initially organized and processed in the olfactory bulb’s Glomerular Layer (31). The exploratory behavior observed in the inner plexiform layer (IPL) and outer plexiform layer (EPL) (GO:0035641 and GO:0035640) may be related to the integration and processing of olfactory information within these regions. Mammals associate odors with punishment or reward so that odors can trigger behaviors(32).

### Pathways related to cancer edge spots

#### Study on PDAC data

On human pancreatic ductal adenocarcinoma (PDAC) data (18), we classify each spot as an edge spot or inner spot was determined based on the whether its neighboring spots belong to the same cell type. Figure 4(b) depicts this distinction, with inner spots colored red and edge spots colored blue. Table 2 presents the top Gene Ontology (GO) terms identified between inner- and edge-cancer cells by our method, highlighting the differences between inner and edge regions of the tumor. Heatmaps of some representative genes associated with the top GO terms (GO:0042742 and GO:0007276) are illustrated in Figure 4(d).

**Table 2.** Top GO terms differentiating edge and inner cancer spots in PDAC data.

| GOID | Term | dCor |
| --- | --- | --- |
| GO:0042742 | defense response to bacterium | 0 |
| GO:0007276 | gamete generation | 0 |
| GO:0034599 | cellular response to oxidative stress | 0 |
| GO:0006816 | calcium ion transport | 0 |
| GO:0071559 | response to transforming growth factor beta | 0 |
| GO:0002443 | leukocyte mediated immunity | 0.083 |
| GO:0046034 | ATP metabolic process | 0.119 |
| GO:0042327 | positive regulation of phosphorylation | 0.125 |
| GO:0046700 | heterocycle catabolic process | 0.142 |
| GO:0098916 | anterograde trans-synaptic signaling | 0.162 |

The identified Gene Ontology (GO) terms are associated with the development and progression of cancer, elucidating the complex behavior of cancer cells in environmental adaptation, signal transduction, cell migration, and immune evasion. One notable term is calcium ion transport, which involves TRPM7’s regulation of Ca2+ levels, impacting calcium-permeable ion channels. This regulation can modify signaling pathways essential for survival, cell cycle progression, proliferation, growth, migration, invasion, and epithelial-mesenchymal transition (EMT), thereby influencing cell behavior and fostering tumor growth. Enhanced calcium ion transport around the tumor may promote cancer cell migration and invasion (33). Other identified terms include chemosynaptic transmission and trans-synaptic signaling. Researches have demonstrated that dynamic interactions between cancer cells and neurons can enhance the invasiveness of PDAC, thereby supporting tumor occurrence, growth, and invasion (34). Additionally, regulation of cell growth is also identified. TGF-*β*, a critical growth factor, plays a significant role in regulating cell proliferation, migration, and differentiation. During cancer progression, tumor cells may overcome the inhibitory effects of TGF-*β* signaling through mutations, and TGF-*β* can induce EMT, resulting in the loss of cell polarity and increased invasiveness of cancer cell (35). Furthermore, Cancer cells at the tumor margin may undergo oxidative stress and cell death (36), which explains the identification of cellular responses to oxidative stress.

#### Study on OSCC data

We apply our method to the spatial transcriptomic data of HPV-negative Oral Squamous Cell Carcinoma data (OSCC)(37), measured using 10xGenomics Visium platform. The image of H&E-stained is shown in Figure 4(d), with detailed annotations of the regions in Figure 4(e). These areas include Squamous Cell Carcinoma (SCC), Lymphocyte Negative Stroma, Lymphocyte Positive Stroma, and Muscle. Metastasis and invasion are the major causes of mortality in OSCC patients (38), so understanding the microenvironmental functions at the tumor edge area is particularly important. Our focus is on analyzing SCC tumor cells to investigate if there are significant biological function differences between the inner area and edge area. The study area is highlighted in red in Figure 4(d). Tumor spots are divided into inner and edge based on the type of surrounding cells, with inner spots in orange and edge spots in blue, as shown in Figure 4(f). Table 3 presents the top four pathways with the most notable differences.

**Table 3.** Top GO terms differentiating edge and inner cancer spots (OSCC).

| GOID | Term | dCor |
| --- | --- | --- |
| GO:0007186 | G protein-coupled receptor signaling pathway | 0.113 |
| GO:0071496 | cellular response to external stimulus | 0.248 |
| GO:0010975 | regulation of neuron projection development | 0.276 |
| GO:0048545 | response to steroid hormone | 0.291 |

A critical pathway identified is the G protein-coupled receptor signaling (GPCRs) pathway (GO:0007186), which can regulate cell proliferation, activity, and movement. The signal pathway is key to tumor growth, angiogenesis, and metastasis. At the interface between the tumor and lymphocyte-negative stroma, malignant cells may utilize the normal function of GPCRs to evade immune surveillance, enhance nutrition and oxygenation, and infiltrate surrounding tissues (39, 40). The cellular response to external stimuli (GO:0071496) includes reactions to various external factors, such as increased interstitial pressure, changes in enzymatic activities, and pH level shifts. An increase in invasiveness and metabolic activity within tumor tissues leads to localized hypoxia in the tumor microenvironment, thereby raising the levels of regulatory transcription factors, such as hypoxia-inducible factors (HIFs) at the tumor edges. This hypoxic stimulus might further inhibit central metabolic synthesis, enhancing invasiveness and anti-apoptotic capabilities. At the edges, tumor cells may need to respond more to stimuli from the immune system, including cytokines released by immune cells (41–43). The regulation of neurite outgrowth in oncology might relate to the proliferation of perineural fibers around tumors, which could be particularly prominent at tumor edges (related to GO:0010975, regulation of neuron projection development). Perineural invasion includes direct tumor invasion into surrounding nerve sheaths and the secretion of neurotrophic factors by tumor cells to promote cancer cell invasion and neurite growth (44). Up to 80% of OSCC exhibits perineural infiltration, often resulting in pain or even cranial neuropathies (45). Additionally, the response to steroid hormone signals (GO:0048545), such as estrogens, could enhance the motility of cancer cells by participating in the epithelial-mesenchymal transition (EMT) process, crucial for the tumor’s invasiveness, dissemination, and metastasis capabilities. Evidence suggests that OSCC cell proliferation and invasiveness significantly increase following *β*-estradiol stimulation (46).

## Discussion

In this work, we developed two related spatial pathway testing methods that by-passes the selection of spatially variable genes, as well as allows nonlinear and complex relations. Both are based on the testing of statistical dependencies between random vectors. One method is to find spatially associated pathways in an unsupervised manner, and other to find pathways associated with predetermined regions in a supervised manner. They can generate easily interpretable biological results from spatial transcriptomics data. We believe they are valuable addition to the tools for the analysis of spatial transcriptomics data.

## Data availability

Our study use publicly available datasets. Human PDAC dataset (GSE111672) available for download at human PDAC dataset (https://www.ncbi.nlm.nih.gov/geo/query/acc.cgi?acc=GSE111672).

Adult mouse hippocampus (GSM4656181) is publicly available in the National Center for Biotechnology Information (NCBI) repository (https://www.ncbi.nlm.nih.gov/geo/query/acc.cgi?acc=GSM4656181). Oral squamous cell carcinoma (GSE208253) is publicly available in the National Center for Biotechnology Information (NCBI) repository (https://www.ncbi.nlm.nih.gov/geo/query/acc.cgi?acc=GSE208253).

## Funding

This work was partially supported by the National Key R&D Program of China (2022ZD0116004), Guangdong Talent Program (2021CX02Y145), Guangdong Provincial Key Laboratory of Big Data Computing, and Shenzhen Key Laboratory of Cross-Modal Cognitive Computing (ZDSYS20230626091302006).

## Notes

### Competing Interest Statement

The authors have declared no competing interest.

